# A Pan-Cancer Transcriptome Analysis Reveals Pervasive Regulation through Tumor-Associated Alternative Promoters

**DOI:** 10.1101/176487

**Authors:** Deniz Demircioğlu, Martin Kindermans, Tannistha Nandi, Engin Cukuroglu, Claudia Calabrese, Nuno A. Fonseca, Andre Kahles, Kjong Lehmann, Oliver Stegle, Alvis Brazma, Angela N. Brooks, Gunnar Rätsch, Patrick Tan, Jonathan Göke, on behalf of the PCAWG Transcriptome Working Group, and the ICGC/TCGA Pan-Cancer Analysis of Whole Genomes Network

## Abstract

Most human protein-coding genes are regulated by multiple, distinct promoters, suggesting that the choice of promoter is as important as its level of transcriptional activity. While the role of promoters as driver elements in cancer has been recognized, the contribution of alternative promoters to regulation of the cancer transcriptome remains largely unexplored. Here we infer active promoters using RNA-Seq data from 1,188 cancer samples with matched whole genome sequencing data. We find that alternative promoters are a major contributor to context-specific regulation of isoform expression and that alternative promoters are frequently deregulated in cancer, affecting known cancer-genes and novel candidates. Our study suggests that a highly dynamic landscape of active promoters shapes the cancer transcriptome, opening many opportunities to further explore the interplay of regulatory mechanism and noncoding somatic mutations with transcriptional aberrations in cancer.

## INTRODUCTION

The key element in regulation of transcription is the region upstream of the transcription start sites (TSS), the promoter. Promoters contain the elements required to initiate transcription, and they integrate the signals from distal regulatory elements and epigenetic modifications that together determine the level of transcription. In the human genome, the majority of protein coding genes are regulated by multiple promoters that initiate transcription for different gene isoforms (Carninci et al., 2006; Sandelin et al., 2007). In contrast to alternative splicing which regulates gene isoform expression post-transcriptionally, the usage of alternative transcription start sites provides a way to regulate gene isoform expression pre-transcriptionally (Ayoubi & Van De Ven, 1996). Therefore, promoters not only determine when a gene is active and how active it is, they also regulate which gene isoform will be expressed.

In cancer, somatic mutations, genomic re-arrangements, and changes in the regulatory or epigenetic landscape have been found to affect the promoter of several oncogenes, and it has been suggested that promoters contribute to the malignant transformation of the cells (Khurana et al., 2016; Sharma, Kelly, & Jones, 2010; Vogelstein et al., 2013). Genome-wide studies of promoters using the H3K4me3 histone modification, an epigenetic mark found at active promoters, or CAGE tag sequencing of the 5’ end of transcripts have found that transcription start sites frequently are differentially used in cancer (Bernstein et al., 2002; Chi, Allis, & Wang, 2010; Gherardi, Birchmeier, Birchmeier, & Vande Woude, 2012; Hashimoto et al., 2015; Kaczkowski et al., 2016; Kodzius et al., 2006; Muratani et al., 2014; Santos-Rosa et al., 2002; Takahashi, Kato, Murata, & Carninci, 2012). However, as data such as H3K4me3 profiles or CAGE-Tag is not available for most cancer studies, the role of alternative promoters in cancer remains largely unexplored.

Because any change in a cell’s identity and function will be reflected in a change in gene expression, transcriptome profiling by RNA-Sequencing is one of the most widely studied large-scale molecular phenotypes in cancer. Analysis of gene expression in cancer has uncovered fundamental insights of tumor biology (Cancer Genome Atlas, 2012), enabled stratification of cancer types (Cancer Genome Atlas Research, 2012), predicted clinical outcome (Gerstung et al., 2015), and guided treatment decisions (Cancer Genome Atlas Research, 2011), forming a cornerstone of data driven precision oncology. RNA-Seq data measures the transcriptome largely unbiased, and as promoters regulate expression of isoforms with distinct 5’ start sites, it could potentially be used to identify active promoters (Feng et al., 2016; Pal et al., 2011; Reyes & Huber, 2017).

In this manuscript, we infer active promoters from RNA-Seq data, enabling the analysis of promoter activity in thousands of samples using publicly available expression data, thereby generating the largest currently available catalog of active promoters in human tissues and cancers. We apply this approach to comprehensively analyze alternative promoters in the Pan-Cancer-Analysis of Whole Genomes (PCAWG) cohort of 1188 patient samples with matched whole genome sequencing data covering 27 different cancer types (Calabrese et al., 2018), and we compare promoter usage to more than 1,500 normal tissue samples (The GTEx Consortium, 2017). We find that alternative promoters are frequently used to increase isoform diversity and that a large number of known cancer genes and novel candidates show deregulation of promoters in cancer. Our data suggests that the landscape of active promoter is highly dynamic and associated with patterns of somatic mutations, and that patient-to-patient variation in promoter activity is associated with survival. We propose that the precise knowledge of which promoter is active in each patient helps understanding the genetic, transcriptional, and pathological profile of individual tumors.

## RESULTS

### Identification of active promoters in 1,209 cancer samples from 27 cancer types

The promoter is defined as the regulatory region upstream of the transcription start site. Using the Ensembl v75 annotations (Yates et al., 2016), we compiled a set of 112,985 possible promoters, assuming that isoforms which have identical or very close TSSs are regulated by the same promoter (Frith et al., 2008). We then define promoter activity as the total amount of transcription initiated at each promoter. By quantifying the expression that is initiated at each promoter we can then infer levels of promoter activity from RNA-Seq data (Fig. 1a). As the number of promoters is much smaller than the number of isoforms per gene, the problem of promoter activity estimation is heavily reduced in complexity, resulting in more robust inference (Supplementary Fig. 1a). To further reduce the number of false positives, we restrict the analysis to promoters that can be uniquely identified (Supplementary Fig. 1b). Following this approach, we quantified promoter activity in 1,209 samples from the PCAWG cohort covering 27 cancer types. Across all samples we identified the most active promoter (major promoter) for 16,694 genes, we identified 21,313 additional promoters that are active at lower levels (minor promoters), and we found 56% (48,312) of promoters to be inactive (Fig. 1b, Supplementary Fig. 1c). In the absence of regulatory genomics data, the first promoter of a gene is often assumed to be active. Interestingly, our data shows that the dominating major promoters can occur at any position within a gene. We find that 1 out of 3 major promoters are located downstream of the first TSS (Fig. 1c), demonstrating how RNA-Seq data adds information and context to genome annotations.

**Figure 1:**
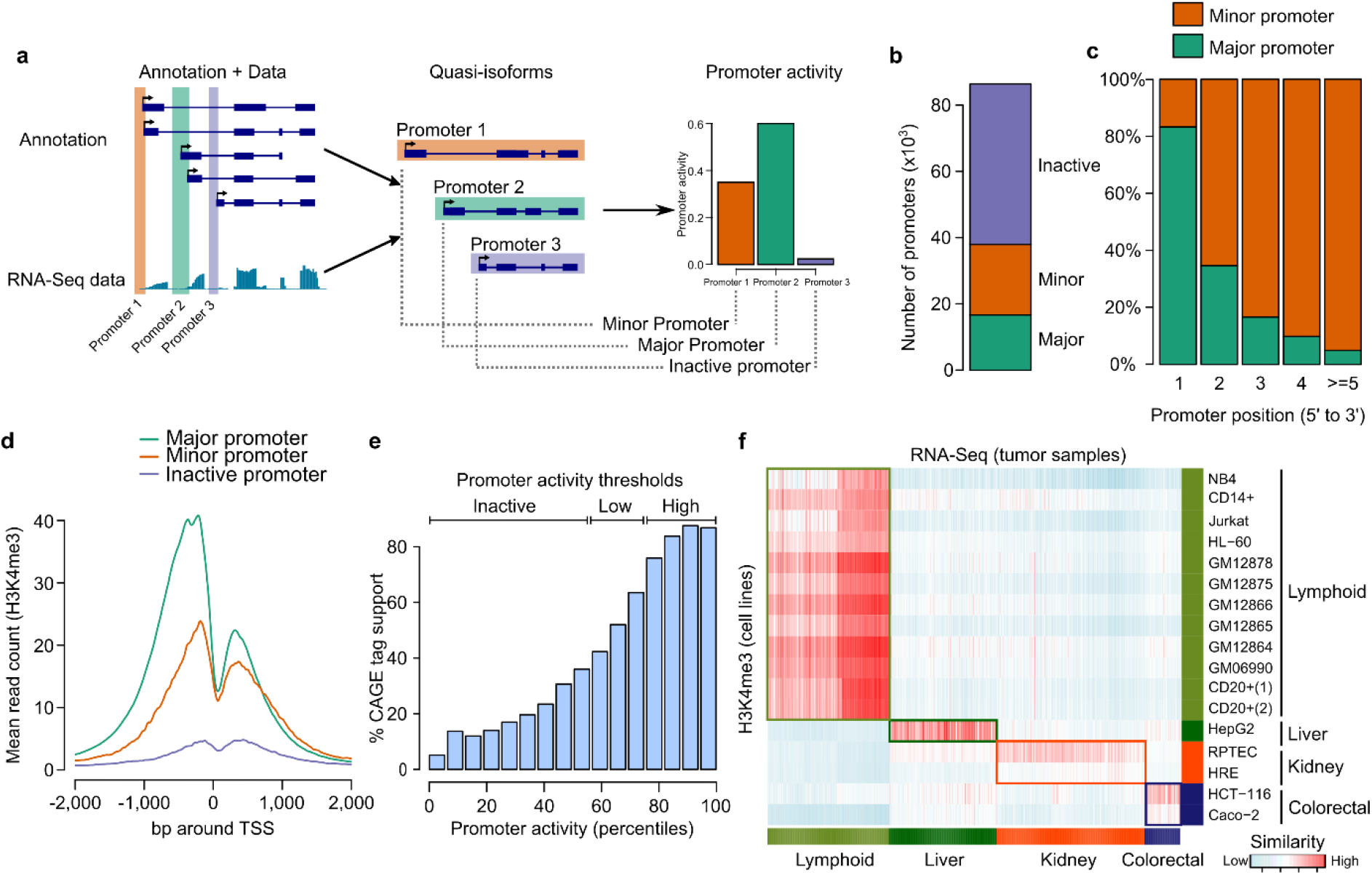
Promoter activity estimation using RNA-Seq data and comparison with ChIP-Seq and CAGE data. (**a**) Schematic representation of promoter activity quantification using RNA-Seq data. Isoforms which are controlled by the same promoter are grouped and promoter activity is estimated as the total expression of these grouped isoforms. (**b**) Categorization of annotated promoters based on the average activity estimate across all samples. Major: most active promoter for a gene; minor: active promoter, lower activity than major promoter; inactive: no activity detected (threshold: 0.5). (**c**) Major/minor promoter proportions across TSSs ranked by position (5’ to 3’), based on multi promoter genes with at least one active promoter. (**d**) Mean H3K4me3 ChIP-Seq signal across 59 ENCODE cell lines for the pancancer major (green), minor (red), and inactive promoters (purple) at the first TSS. (**e**) Percentage CAGE tag support for inactive, low, and high activity promoters. (**f**) Correlation of promoter activity estimates and H3k4me3 ChIP-Seq signal for lymphoid, liver, kidney, and colorectal ENCODE cell lines and PCAWG samples. PCAWG samples show higher correlation with ChIP-Seq data from the matching tissue.

To evaluate the accuracy of expression-based estimation of promoter activity, we compared them to publicly available H3K4me3 ChIP-Seq and CAGE tag data from a variety of different cell lines and tissues (The ENCODE Consortium, 2012; The FANTOM Consortium, 2014; Lizio et al., 2015). The major promoters identified in the PCAWG cohort show the highest levels of H3K4me3 and CAGE tag support, whereas promoters identified as inactive show the lowest H3K4me3 levels and CAGE tag support, demonstrating that expression and epigenetics based estimates show a remarkable level of consistency (Fig. 1d, e, Supplementary Fig. 1d, e, f). Furthermore, estimates from cancers were most similar to ChIP-Seq profiles from matching cell lines (Fig. 1f). Interestingly, while promoter activity estimates from patients were generally highly consistent, cell lines showed a much higher variance (Supplementary Fig. 1g). It has been observed before that cancer cell lines differ from the primary tissue (The FANTOM Consortium, 2014), suggesting that RNA based estimates more accurately reflect the promoter landscape of the tumor than cell line based estimates. Overall, this analysis demonstrates that RNA-Seq data enables the quantitative, robust, and reproducible estimation of promoter activity.

### Alternative promoters are a major contributor to isoform diversity

Genome-wide, we find that promoter activity is dominated by the tissue and cell of origin for each cancer type (Fig. 2a). This closely resembles the observation from gene expression, despite using only the minimal set of discriminative reads indicative of promoter activity (Supplementary Fig. 2a). In contrast to gene-level expression estimates, promoter activity enables us to investigate the contribution of each promoter to the overall expression pattern. Among all expressed protein-coding genes, 49% have at least 2 active promoters that contribute to more than 10% of the overall gene expression (Fig. 2b, c). In principle, these promoters are independent regulatory units which can be used in a different context to control changes in isoform expression. The usage of such *alternative promoters*-promoters whose activity depends on the context but not on the activity of the gene’s remaining promoters - will not be detectable with gene level based expressions analysis. Therefore, even though globally promoter activity reflects gene expression, there is additional information in promoter activity that cannot be detected at the gene expression level.

**Figure 2:**
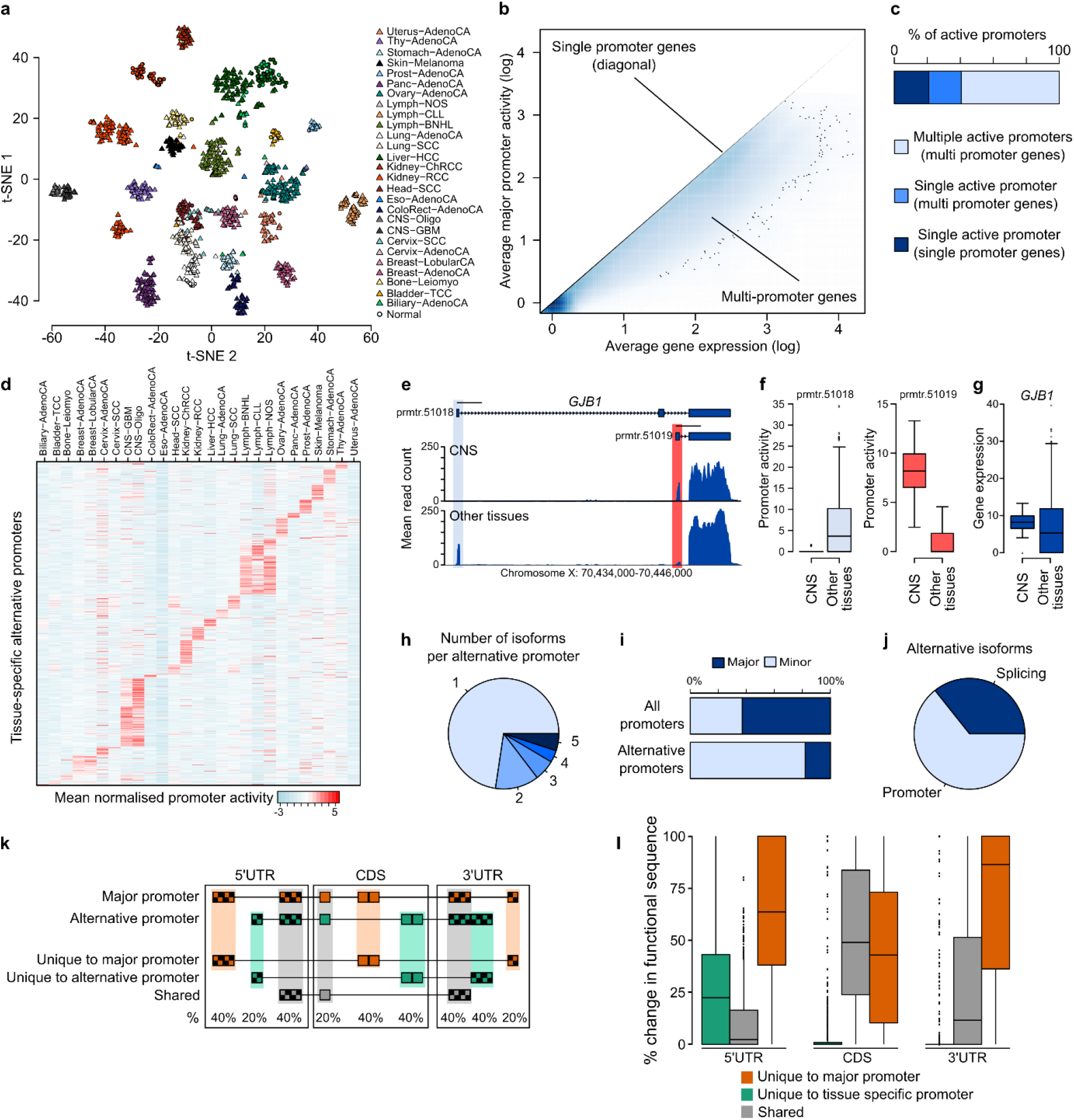
Alternative promoters are a major contributor to isoform diversity. (**a**) t-SNE plot using the top 1,500 promoters with the highest variance in promoter activity. (**b**) Comparison of major promoter activity and gene expression (sum of all promoters). A single promoter often does not fully explain gene expression, minor promoters contain additional information. (**c**) Most active promoters are observed at genes with multiple promoters. (**d**) Heatmap showing the normalized mean promoter activity for tissue-specific alternative promoters of genes which do not change in overall expression. (**e**) Shown is the mean read count at the *GJB1* gene locus for samples from the central nervous system (CNS) and from all other tissues. The light blue and red regions highlight the two alternative promoters. (**f**) The second *GJB1* promoter (prmtr.51019) is more active in CNS compared to all other tissues, whereas the first promoter (prmtr.51018) is inactive in CNS samples. (**g**) Comparison of gene expression levels for *GJB1*.(**h**) Number of isoforms that can be transcribed from tissue-specific alternative promoters. (**i**) Alternative promoters are most often minor promoters. (**j**) Shown is the fraction of isoform switching events across tissues that involve a change in promoter. (**k**) Schematic representation for the analysis of 5’UTR, CDS, and 3’ UTR regions and their association with alternative promoter usage. Regions unique to the major and alternative promoter, and regions shared among them are quantified for each tissue-specific alternative promoter. (**l**) Shown is the percentage of the 5’UTR, 3’UTR, and CDS sequence that is unique to the tissue specific promoter gene isoforms (green), unique to the major promoter gene isoforms (blue), and that is shared (gray). Tissue-specific promoters shorten the coding sequence by 40% on average.

To approximate the prevalence of alternative promoters as context-specific regulators of transcription, we searched for promoters that show significantly changed activity across tissues at genes that do not show an overall change in expression (Fig. 2d, Supplementary Fig. 2b, c). Strikingly, our data demonstrates that even genes that do not show any tissue-specificity at the gene expression level can be under control of 2 independent, highly tissue-specific alternative promoters which regulate distinct gene isoforms (Fig. 2e, f, g). The majority of tissue-specific alternative promoters activate single isoforms, providing a direct link between transcriptional regulation and isoform expression (Fig. 2h). Alternative promoters often correspond to minor promoters that are expressed at lower levels compared to the constitutively active major promoter (Fig. 2i). However, for 18% of genes we observe that the major promoter is switched (Fig. 2i). Interestingly, on a global level, 60% of all isoform switching events involve a switch in promoters (Fig. 2j), demonstrating that alternative promoters are a major contributor to tissue-specific transcriptional diversity.

To understand the consequence of alternative promoter usage on the gene product, we examined how the functional regions (5’ UTR, CDS, 3’UTR) differ compared to the major promoter (Fig. 2k). As expected, use of an alternative promoter is almost always associated with a change in the 5’ UTR region (Supplementary Fig. 2d), with on average less than 20% of the 5’UTR sequence being shared between alternative promoters (Fig. 2l). A change in promoters also dramatically effects the coding part of RNAs, often involving a change of almost 50% of the protein coding sequence (Fig. 2l). Surprisingly 90% of alternative promoters encode for isoforms that potentially use a different 3’ UTR sequence based on annotations (Supplementary Fig. 2d). This suggests that promoters not only regulate transcription initiation, but that they specifically regulate alternative isoforms that are marked by distinct sequences, possibly influencing post transcriptional regulation, translation, and protein structure in a context-specific manner.

### Cancer-associated promoters regulate isoform switching of oncogenes and tumor suppressors

Many cancer-associated genes and pathways have been discovered by comparing the expression profile of cancer with the expression profile of normal tissues (Fay et al., 2003; Gross, Kreisberg, & Ideker, 2015; Hippo et al., 2002; Rapin et al., 2014). The large number of context-specific alternative promoters found in this study suggests that promoters might be among the unknown driving forces behind the transcriptional changes in cancer. To investigate this hypothesis, we searched for promoters that show a change in activity in cancer compared to normal tissue using adjacent samples from the PCAWG data set and additionally 1,831 samples from GTEx (v6.p1) (The GTEx Consortium, 2017) (Fig. 3a). For the majority of tumor types the most similar tissue is indeed the tumor tissue (Fig. 3b, Supplementary Fig. 3a). Interestingly, lung squamous cell carcinomas and bladder carcinomas are most similar to normal skin tissue, reflecting the cell or origin for these tumors (Cancer Genome Atlas Research, 2014). Using these matched tissue groups, we then identified cancer-associated alternative promoters. For each tissue we find between 93 and 226 promoters that are differentially regulated in cancer compared to normal (Fig. 3c, Supplementary Fig. 3b, d). The change in expression due to cancer-associated promoters is largely independent from the other promoters for each gene, confirming that alternative promoters indeed act as independent regulatory units which can specifically be deregulated in cancer (Fig. 3d, Supplementary Fig. 3c, e). Our analysis recapitulates promoter switching events that have been associated with cancer, amongst others for *MEST* (Fig. 3e, f) (Nakanishi et al., 2004) and *MET* (Gherardi et al., 2012). However, the vast majority of events are novel and have not been described before. Among the genes that show alternative promoter activation in cancer are known cancer biomarkers such as *SEPT9* (deVos et al., 2009) or *TNFRSF19 (TROY)* (Paulino et al., 2010), oncogenes and tumor suppressors such as *IKZF1* (Boer et al., 2016) which has been reported to be involved in human B-cell acute lymphoblastic leukemia, the well described protooncogene *CTNNB1* (β-catenin) (Lazar et al., 2008), *BID* (Supplementary Fig. 3f) (Lee et al., 2004), a pro-apoptotic target gene of p53, or *MLLT1* (Perlman et al., 2015), which has been associated with childhood kidney cancer (Supplementary Fig. 3g). To understand the underlying regulatory changes leading to the use of cancer associated promoters, we performed a de-novo motif analysis. Across all cancer types we find several enriched transcription factor motifs (Supplementary Fig. 3l-q), suggesting that changes in the activity of promoters are partially driven through a change in the regulatory networks.

**Figure 3:**
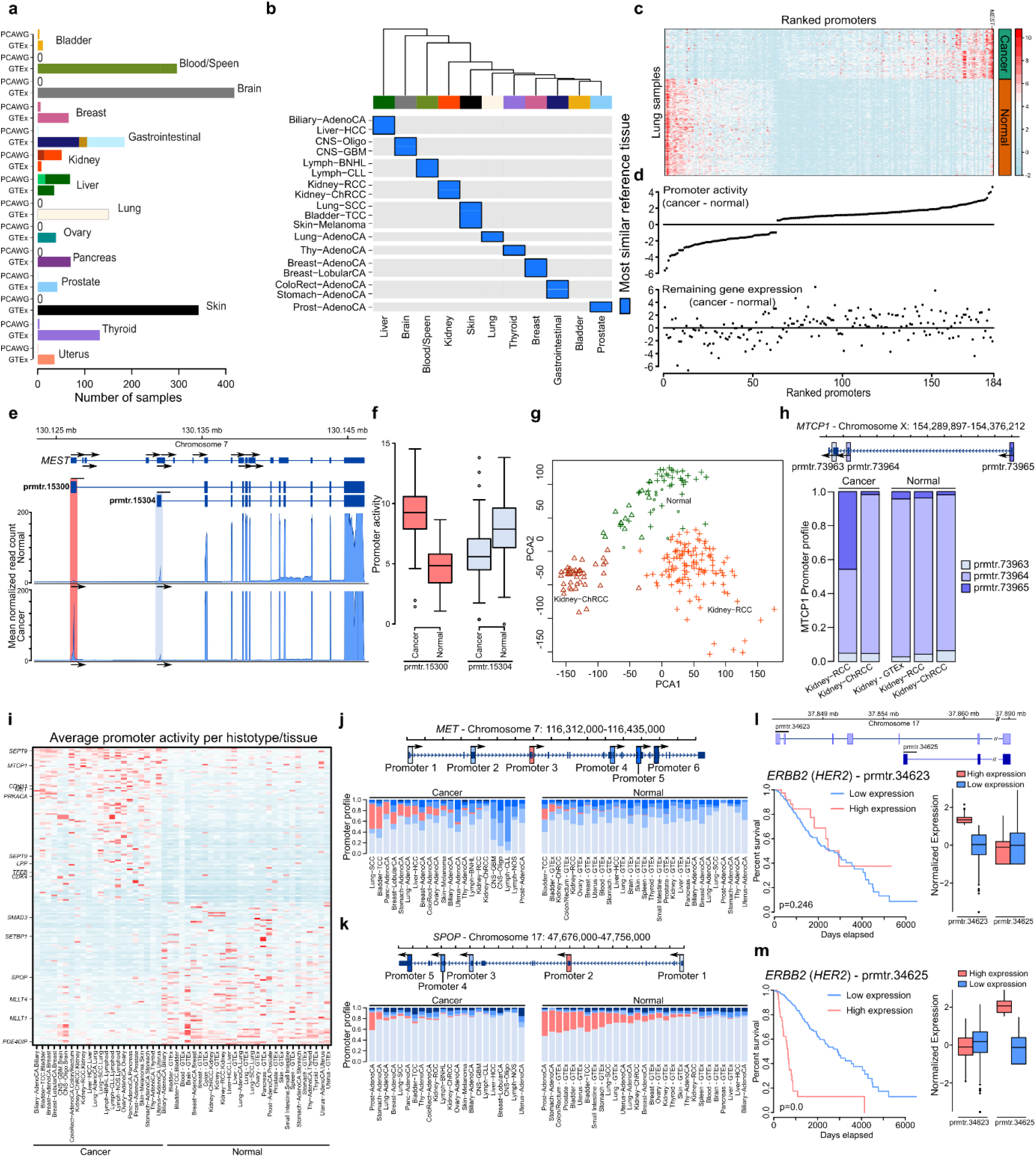
Cancer-associated alternative promoters regulate oncogenes and tumor suppressors both for individual tissue types and across cancers. (**a**) Overview of cancer and normal data obtained by combining PCAWG and GTEx samples. (**b**) Cancer samples are most similar (highest average correlation of promoter activity) to the normal samples from the same tissue type. Tumor types with less than 15 normal and cancer samples are excluded from this analysis, normal samples are batch corrected to account for different data sources (PCAWG and GTEx). (**c**) Heatmap showing the normalized promoter activity estimates for lung cancer and normal samples, ranked by mean difference. (**d**) Difference in cancer-associated promoter activities (upper panel) and gene expression excluding alternative promoters (lower panel) for lung cancer and normal samples, ranking is similar to heatmap in (**c**). (**e**) Shown is the mean read count at the *MEST* gene locus for lung cancer and lung normal samples (left panel). The light blue and red regions highlight the cancer-associated promoters. First promoter is the major promoter in cancer samples whereas second promoter is the major promoter in the normal samples. (**f**) The promoter activity for the first (red) and second (light blue) promoters of *MEST*in cancer and normal samples show the switch of promoters between cancer and normal samples. (**g**) PCA plot using the top 1,000 promoters with the highest variance in promoter activity in kidney samples. (**h**) Shown is the *MTCP1* locus (top) and mean relative activity across different kidney cancer and normal samples (bottom). The first promoter of *MTCP1*(prmtr.73965, dark blue) displays Kidney-RCC tumor subtype specific activation. (**i**) Heatmap showing the average activity of promoters that significantly differ between cancer and normal samples across multiple cancer types (pan-cancer associated promoters). Promoters of known cancer-associated genes are highlighted (Futreal et al., 2004). (**j, k**) Relative activity profile of pan-cancer-associated promoters. Shown is the *MET* gene locus which shows activation of an alternative promoter in cancer (**j**); and the *SPOP* gene locus which shows lower activity of an alternative promoter across multiple cancer types (**k**). (**l, m**) Shown is the *ERBB2* gene locus (top). Increased activity of the second (minor) promoter (prmtr.34625) is associated with decreased survival in lower grade glioma patients (**m**), whereas increased expression of the first (major) promoter (prmtr.34623) is not associated with survival (**l**). The boxplots on the right panels show the activity of each promoter in their respective high (red) and low (blue) activity sample groups i.e. the same sample groups as the survival plots on the left panels.

Interestingly, alternative promoters also differ between closely related tumor types from the same tissue. For the 2 different kidney tumor types we find a large number of genes that only show minor changes in overall gene expression, but where alternative promoter usage causes a significant, tumor-type-specific change in isoform expression (Fig. 3g, 3h, Supplementary Fig. 3h, i). We have also identified several genes, such as *STAU2*, that use distinct alternative promoters across the clinical subtypes of breast cancer (Supplementary Fig. 3r, s, t). While some promoters were specifically deregulated in single tumor types, we observed that a number of alternative promoters were deregulated in multiple tumor types from different tissues compared to their matched normal counterpart. Overall, we find 204 such promoters, several of which belong to known oncogenes and tumor suppressors (Futreal et al., 2004) (Fig. 3i). Among the known cancer-associated genes for which we find differentially activated promoters are the previously described *MET* gene (Fig. 3j), and genes for which usage of an alternative promoter in cancer has not been described such as *LSP1*, or *SPOP* (Fig. 3k). Again, we find that the choice of promoters changes the 5’UTR, CDS, and 3’UTR sequences, indicating that transcriptional changes in cancer are translated into functional differences in the gene product (Supplementary Fig. 3j, k). Together, our analysis highlights that alternative promoter usage provides a major source of transcript diversity affecting known cancer genes and new candidates, demonstrating that promoters are a key contributor to the deregulated cancer transcriptome, often independently from an overall change in gene expression.

### Alternative promoter usage of *HER2* is associated with survival

Gene expression has been used as a biomarker to predict cancer patient survival (Director’s Challenge Consortium for the Molecular Classification of Lung et al., 2008; Finak et al., 2008; Salazar et al., 2011). As our data suggests that alternative promoters are often independently regulated, we hypothesized that promoter activity in certain cases might provide a more accurate predictor for genes that use multiple promoters. To test this hypothesis, we investigated the association of promoter activity with survival estimates using TCGA data for cancer census genes that use alternative promoters in cancer. Indeed, we find a number of genes that show a significant association with survival for a specific promoter. Most strikingly, we find that a minor promoter of *ERBB2* (also known as *HER2*) in lower grade glioma patients is predictive of poor outcome, whereas the major promoter shows no significant association with patient survival (Fig. 3l, m). High gene expression levels of *ERBB2* have been associated with aggressive tumor types (Slamon et al., 1987). Our data suggests that such associations can be promoter specific, indicating that survival is either associated with the underlying regulatory changes or with the differential usage of gene isoforms determined through the choice of promoters. Thus, the level of promoter activity can potentially be more specific as a biomarker compared to gene expression, demonstrating the promise to further explore their role in cancer.

### Patterns of noncoding promoter mutations in cancer

Accumulation of somatic mutations plays a central role in cancer not only by affecting protein coding genes, also by disrupting noncoding gene regulatory elements (Kandoth et al., 2013; Rheinbay, Parasuraman, et al., 2017; Weinhold, Jacobsen, Schultz, Sander, & Lee, 2014). It was found that mutational patterns are highly heterogeneous (Lawrence et al., 2013; Maruvka et al., 2017). To better understand which properties of promoters are associated with accumulation of somatic (single nucleotide) mutations in cancer, we investigated the whole genome sequencing data for all patients with matched RNA-Seq data in the PCAWG cohort. We find that promoters of genes with a less complex promoter architecture show higher number of mutations (Fig. 4a). These genes are more often non-coding (Fig. 4b) and within regions associated with later replication timing (Fig. 4c), confirming that distinct groups of promoters are exposed to different mutational patterns. Besides these genomic properties of promoters, we also find that their activity and the tissue of origin influence the observed mutation burden: major promoters show the highest enrichment of mutations in a window 200bp upstream of the TSS, whereas minor and inactive promoters show lower levels of mutations (Fig. 4d, Supplementary Fig. 4a). In agreement with previous findings (Sabarinathan, Mularoni, Deu-Pons, Gonzalez-Perez, & Lopez-Bigas, 2016), Skin-Melanoma displayed the highest variation of mutation rates (C>T) due to inefficiency of nucleotide excision repair (NER) in fixing lesions induced by UV light when a transcription factor is bound upstream TSS (Supplementary Fig. 4b, c). We observe similar mutational patterns for lung adenocarcinomas and lung squamous cell carcinomas, all of which are cancer types affected by smoking related carcinogens where NER plays a central role in DNA repair (Supplementary Fig. 4a). The cancer type that shows the strongest deviation from this pattern is colorectal adenocarcinoma (Fig. 4e, Supplementary Fig. 4d, e). These data suggest an intricate relation between promoter activation, their genomic properties, and cancer-specific mutational processes that affect the observed mutation burden. Understanding these variations in mutation heterogeneity has implications for identifying driver mutations by determining background mutation rates accurately. A small set of driver promoter mutations (*TERT*, *PAX5*, *WDR74*, *HES1*, *IFI44L*, *RFTN1*, and *POLR3E*) has been identified in the PCAWG cohort, and it is expected that the number increases with higher sample number (Rheinbay, Getz, & PCAWG-2-5-9-14, 2017). As RNA-Seq data is among the most widely generated data, our approach will provide a powerful tool to better understand the relation between somatic mutations, promoter activity and regulatory drivers in cancer.

**Figure 4:**
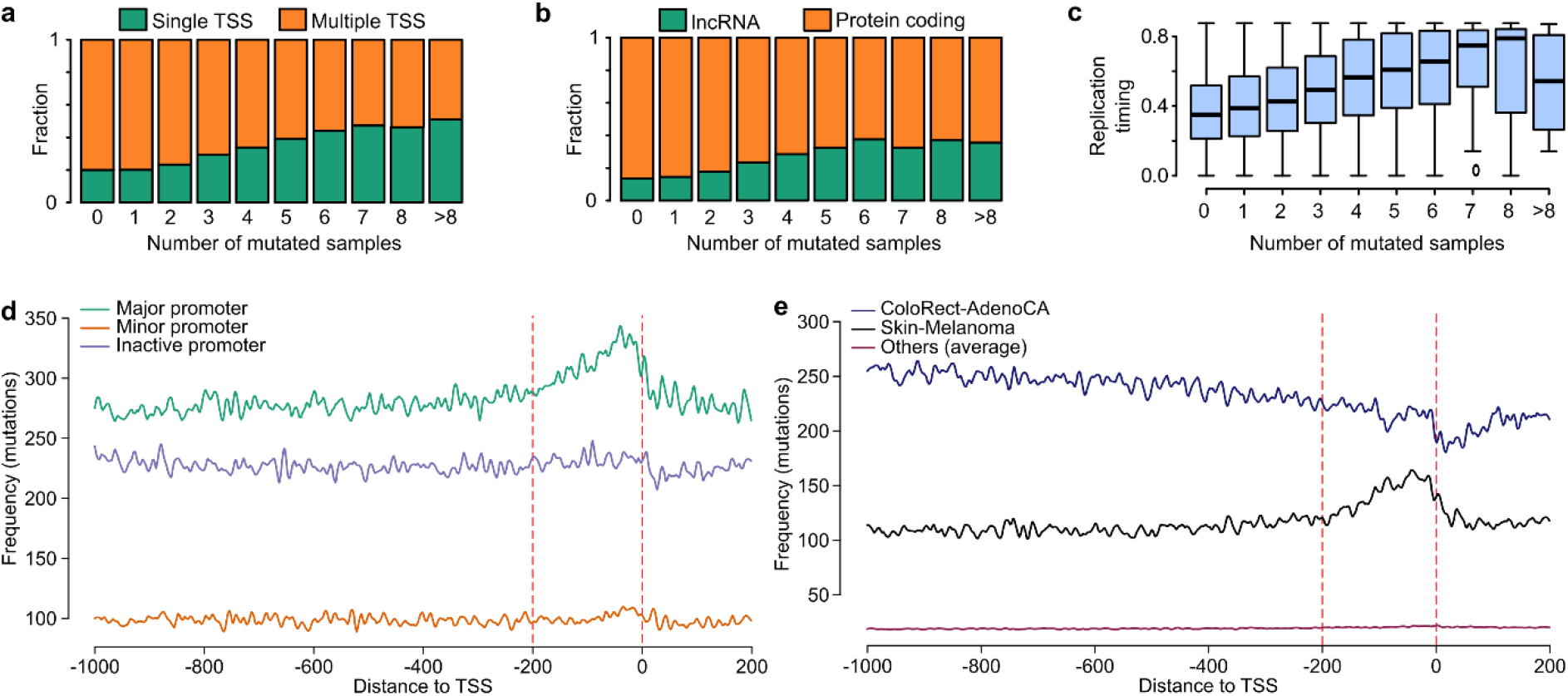
Overview of noncoding promoter mutations patterns across different cancer types and pan-cancer. (a) Proportion of single and multi TSS genes for different numbers of mutations at promoters. Single TSS genes are more frequently mutated than genes with multiple promoters (multiple TSS genes). (**b**) lncRNAs are more frequently mutated at the promoter than protein coding genes. (**c**) Boxplot comparing the replication timing across samples with different number of mutations, a higher number of mutations is associated with later replication timing. (**d**) Distribution of promoter mutations around promoters across PCAWG cohort for major, minor and inactive promoters. Red lines indicate the window of 200bps upstream TSS where major promoters show an enrichment of mutations whereas minor and inactive promoters don’t. (**e**) Distribution of promoter mutations around promoters for the top 2 most mutated cancer types (Skin-Melanoma and ColoRect-AdenoCA) and the others. ColoRect-AdenoCA displays a very different mutational pattern compared to all other cancer types.

## DISCUSSION

Promoters are the key elements that link gene regulation with expression. Studies using ChIP-Seq and CAGE tag data have demonstrated a role of alternative promoters in cancer (Kaczkowski et al., 2016; Muratani et al., 2014; Qamra et al., 2017), yet due to limitations in sample numbers for such technologies, the landscape of active promoters and their variation across cancers and patients has not been described. By analyzing more than 1,200 RNA-Seq samples, we provide the largest survey of promoter activity in human tissues and cancers, confirming known examples, and identifying many novel alternative promoters that are associated with cancer. The scale of these data allows us to describe for the first time patient-to-patient variation in promoter usage. Our analysis suggests that the choice of promoter is tightly regulated, has a significant influence on the cancer transcriptome, and indicates that promoters possibly contribute to the cellular transformation of cancer.

By using RNA-Seq data, our approach enables the analysis of promoter activity in the PCAWG cohort without the need for additional experiments. Similar approaches have been applied to normal tissue expression (Reyes et al. 2018), and embryonic stem cells (Feng et al., 2016). Overall these estimates are highly accurate although we observe an increased uncertainty for some promoters due to use of short read sequencing data. In particular, we find that transcription start sites that lie within internal exons or that overlap with splice acceptor sites are difficult to accurately identify. Information from the 3’ end of transcripts can be used to predict their activity, however this approach heavily depends on accurate annotations and high quality isoform abundance estimates, and a high level of uncertainty remains (Teng et al., 2016). Both CAGE tag data and ChIP-Seq data suggest that these “internal” TSSs are much less used compared to the remaining TSSs, therefore our analysis still captures an accurate and comprehensive view of the promoter landscape in cancer, enabling the analysis of patient-specific promoter activity on a much larger scale compared to other genomic assays.

It is known that alternative promoters contribute to isoform diversity (The FANTOM Consortium, 2014; Reyes & Huber, 2017), yet only few such events have been described in a disease context. In cancer, genes such as *MET* (Muratani et al., 2014), *TP73* (Deyoung & Ellisen, 2007), or *ALK* have been reported to use alternative promoters (Wiesner et al., 2015). By analyzing the role of alternative promoters in this large scale cohort we demonstrate that many more cancer-associated genes use alternative promoters, and that their activity systematically alters the cancer transcriptome across all major cancer types. Our results suggest that transcriptional regulation, possibly involving sequence specific transcription factors and epigenetic modifiers, provides a robust way to pre-transcriptionally determine isoform expression in tumors. The choice of promoter often has an impact on the coding sequence, suggesting that a switch in promoters will alter protein isoforms or result in noncoding transcription. Interestingly, we also observe a frequent change in the 3’UTR sequence that contains regulatory elements such as miRNA binding sites (Lai, 2002), indicating a possible relation between pre- and post-transcriptional regulation. Alternative promoters often show lower levels of activity, and the functional consequence of such transcripts remains to be validated. However, we also find a number of promoter switching events that dramatically change the gene product. Such alternative promoters are frequently found in cancer, most of which are unknown, demonstrating that this aspect has a large potential to be further explored.

In summary, our study demonstrates the pervasive role of alternative promoters in context-specific isoform expression, regulation of isoform diversity, and highlights how patient-to-patient variation in promoter activation is possibly linked to pathological properties in cancer. We provide a comprehensive catalog of active promoters and their expression pattern across 27 cancer types that will be a highly useful resource to understanding the roles of gene regulation and noncoding mutations in cancer. Tissue and cancer-specific promoters could also become highly relevant as sensors and tumor-restricted activators for immunotherapy and the development of novel cancer drugs (Nissim et al., 2017), and it will enable accurate designs of genome wide functional screens (Klann et al., 2017; Marx, 2017). As the vast majority of alternative promoters in cancer has not been described before, our study opens numerous possibilities to explore their contribution to tumor formation, diagnosis, or treatment.

## METHODS

### Promoter activity quantification

In this study, we used Ensembl v75 annotation to determine the set of promoters. We examined the first exon of each TSS and combined the TSS’s with overlapping first exons to obtain the set of promoters. We used these promoters in the downstream analysis to quantify promoter activities. Since a single promoter can be composed of multiple TSSs, we choose the TSS with the highest activity in majority of the samples as the TSS of the promoter. In case of a tie, the 5’ most TSS is chosen as the TSS of the promoter considered.

To quantify the activity of each promoter, we used the split reads aligning into the first intron of each promoter. Reads that connect the first exon with downstream exons were first normalized by reads of splice acceptor sites (indicative of usage of this exon as internal exon, not as first exon). We then standardized the read counts by the observed mean read count in each sample (further referred to as promoter activity estimates). To obtain gene expression estimates, we summed up the activities of each promoter belonging to each gene. We normalized each promoter’s activity by the gene expression to obtain relative promoter activities.

After quantifying the promoter activities for each sample, we divided the promoter set into 3 different categories depending on their activity, namely, major, minor and inactive promoters. We mark the promoters with the highest average activity for each gene across the sample cohort as major promoters. Promoters with average activities smaller or less than 0.5 constitute inactive promoters whereas the other promoters of the gene constitute minor promoters.

### ChIP-Seq analysis

To assess the performance of our promoter activity quantification approach, we compared estimated promoter activities from RNA-Seq data with ChIP-seq data obtained from ENCODE project cell lines. We examined the region spanning 2000 bps upstream and downstream of each promoter for H3K4me3 histone modification signals. We used 59 cell lines from different tissue types that have H3K4me3 data available.

### Alternative promoter analysis

We selected the 1500 promoters and genes with the largest variance across the PCAWG data cohort to demonstrate the tissue specific behavior of promoters and genes alike with a T-SNE plot. The “tsne” R package was used to generate T-SNE plots.

We removed transcription starts of exons that overlapped splice acceptors sites from this analysis as we found that their activity is less reproducibly quantified compared to first exons that do not contain splice acceptor sites. We identified promoters with context dependent activity by comparing the relative activity profiles across different conditions using a t-test. We selected the top 5000 promoters with Benjamini-Hochberg adjusted p-values of less than 0.005 as candidates. We further required each promoter to have at least 2 fold change in promoter activity and less than 1.5 fold change in gene expression across different conditions. To filter for inactive genes, we forced a gene expression threshold of 3 and promoter activity threshold of 2 in both conditions. Finally, we required each candidate gene to have at least 2 active promoters in both conditions. We used this approach to identify the tissue specific alternative promoters by comparing samples for each tumor type against all other samples.

### Identification of isoform switch events

We find the major transcript of each gene in each tumor type using the mean activity across all tumor samples (tumor-specific major transcript). Additionally, we find the major transcript based on the pan-cancer mean activity (pan-cancer major transcript). For each tumor type, we identify the changes in major transcript by comparing tumor specific and pan-cancer major transcripts. A change in major transcript can occur via 2 different mechanisms: either the new tumor specific major transcript is regulated by a different promoter than the pan-cancer major transcript (i.e. a promoter switch event), or the promoter is still the same as the pan-cancer major transcript’s promoter but only the major transcript of this promoter is changed (i.e. a splicing event). For each tumor type, we count the number of major transcript changes for both of these mechanisms. Finally, we sum up the number of times the major transcript change has occurred due to splicing for all tumor types. Similarly, we sum up the number of times major transcript change has occurred due to promoter switch for all tumor types.

### 5’UTR, CDS, and 3’UTR analysis

To understand the functional effect of alternative promoters, we compared the major and alternative promoters for the samples of each tumor type. We determined the major promoters by the mean promoter activity across the samples of the corresponding tumor type. Then, we identify the regions unique to the major promoter, alternative promoter and the regions that are common in both. For each of these regions, we looked at the Ensembl annotations to determine the functional composition, i.e. 5’ untranslated region (5’UTR), exon, coding sequence (CDS) and 3’ untranslated region (3’UTR). We determined for each region not only whether we observe these functional regions, but also how much of total region is observed. To obtain the pan-cancer overview, we considered the set of unique promoter changes occurring across all the tumor types.

### Motif analysis

To uncover the putative transcription factors that might be involved in driving the alternative promoter events, we performed de-novo motif analysis using the RSAT – Metazoa webserver (Thomas-Chollier, Darbo, et al., 2012; Thomas-Chollier, Herrmann, et al., 2012). We provided the 200bs upstream region of the transcription start sites of alternative promoters as target sequence, whereas as the reference set we used the genes with at least 2 active non-internal promoters. We identified the enriched transcription factor motifs by using RSAT Webserver’s comparison tool with the 2018 JASPAR transcription factor DNA-binding preferences database (Khan et al., 2018).

### Identification of tumor specific and pan-cancer cancer-associated promoters

In order to identify cancer associated promoters, we downloaded normal samples from the GTEx project in addition to the normal samples from PCAWG. In total, we obtained 3233 normal samples from GTEx across a wide variety of tissue types and processed the same way as PCAWG samples to obtain promoter activities. We removed the batch effect that might originate from using 2 different data sets by using the “removeBatchEffect” function from ‘limma’ R package. We clustered the combined normal samples by hierarchical clustering where the distance measure was 1 – correlation of non-internal promoter activity. For downstream analysis, we removed internal promoters (see above) and used tumor types with at least 15 normal and tumor samples each for tumor type specific analysis.

For the tumor type specific analysis, we used a generalized linear model to obtain p-values for each promoter based on the relative promoter activity. We adjusted the p-values using the Benjamini-Hochberg method and selected the top 5000 promoters with adjusted p-values less than the background p-value. The background p-value is calculated by using all the promoters without any expression filtering (relative, absolute or fold change). For candidate promoters, we enforced at least 2 fold promoter activity change and at most 1.5 fold gene expression change. Additionally, we required each gene to have at least 2 active promoters (absolute promoter activity greater than 0.5 and relative activity greater than 0.1) and at least a gene expression of 3.

Similar to the tumor specific analysis, we examined the promoters that show context dependent activity pan-cancer wide. We used a generalized linear model with cancer/normal states as explanatory variables and used the same expression based filters to remove false candidates. The generalized linear model analysis is performed by using the ‘limma’ R package

### Mutation burden analysis

To calculate the noncoding mutation burden at each promoters, we considered only the single nucleotide variants (SNVs) for donors with available RNA-Seq data and removed the SNVs located at exons of each gene (Synapse ID: syn7364923). Then, for each sample, we counted the number of noncoding SNVs falling in the 200bp window upstream the TSS of each promoter as the mutation burden.

## ACKNOWLEDGEMENT

This work is funded by the Agency for Science, Technology and Research (A*STAR), Singapore. The Genotype-Tissue Expression (GTEx) Project was supported by the Common Fund of the Office of the Director of the National Institutes of Health, and by NCI, NHGRI, NHLBI, NIDA, NIMH, and NINDS. The data used for the analyses described in this manuscript were obtained from the dbGaP accession number phs000424.v6.p1 on 04/09/2015.

---

**Supplementary Figure 1:**
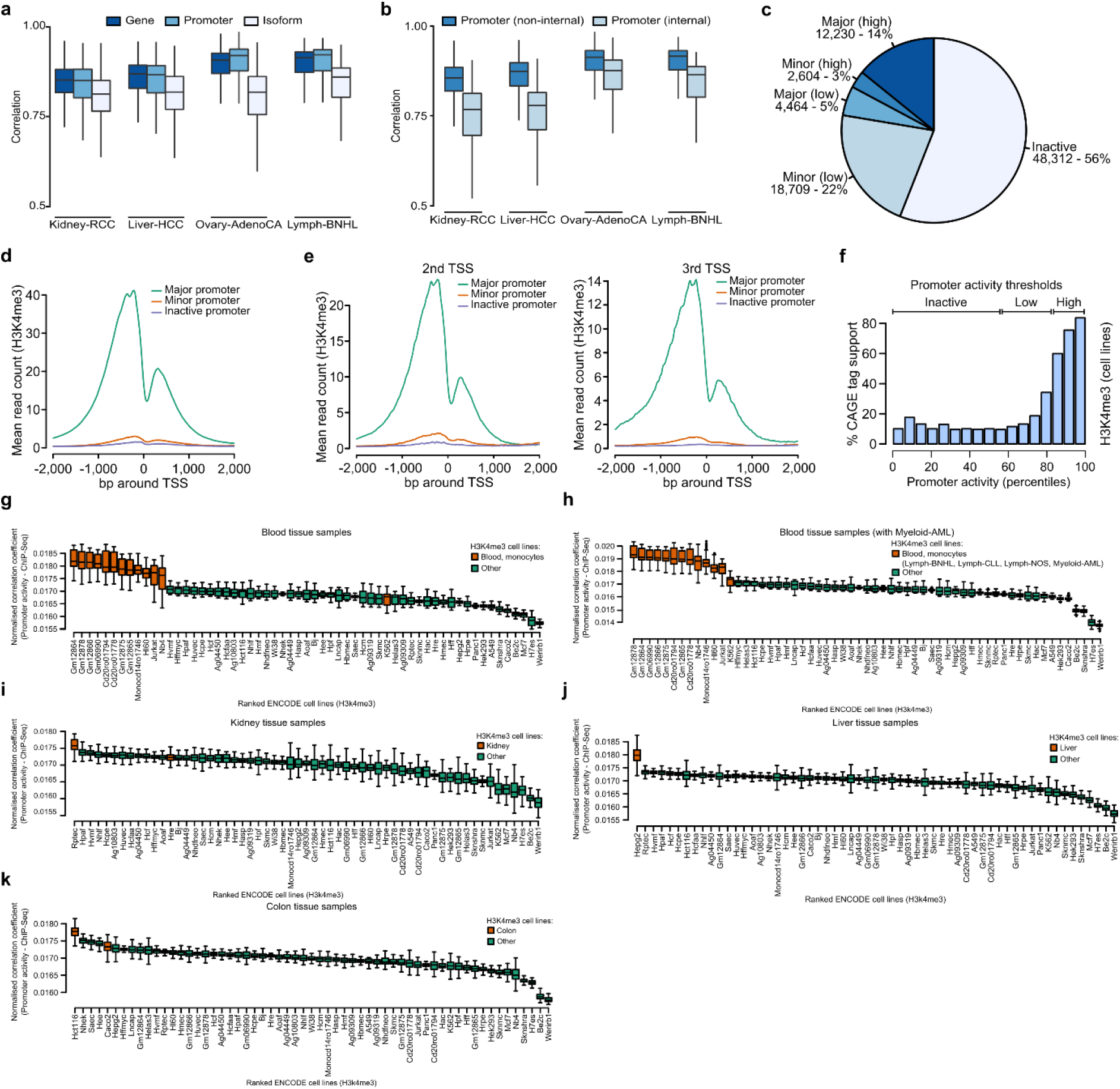
RNA-Seq data can be used to identify active promoters. (**a**) Correlation of expression estimates across samples of the same tumor type for genes (dark blue), active promoters (blue), and isoforms of multi isoform genes (light blue). The correlations are shown for the tumor types with more than 100 samples. A higher correlation of promoter activity estimates suggest a higher level of robustness compared to isoform estimates. (**b**) Correlation of activity for promoters that overlap internal exons (light blue), and promoters that do not overlap internal exons (blue) across the sample pairs of the same tumor type. (**c**) Overview of major (highly expressed), major (lowly expressed), minor (highly expressed), minor (lowly expressed) and inactive promoters across the PCAWG data cohort, using a threshold of 0.5 for low expression, and a threshold of 2.5 to define high expression. (d, e) Mean H3K4me3 ChIP-Seq read count across 59 ENCODE cell lines for the pan-cancer major, minor, and inactive promoters. (**d**) All promoters. (**e**) Promoters at the 2^nd^ and 3^rd^ TSS positions. (**f**) Percentage CAGE tag support for inactive, low, and high activity promoters, shown are only promoters that overlap with internal exons. The CAGE tag support is lower compared to non-internal promoters. (**g - k**) Correlation of promoter activity estimations with H3K4me3 ChIP-Seq read counts from ENCODE cell lines for blood (**g, h**), kidney (**i**), liver (**j**), colon (**k**) tissues. Orange color denotes the matching cell lines in terms of tissue origin. Inclusion of the gray-listed Myeloid-AML samples into promoter activity estimates increases the similarity to the K562 cell line, which otherwise is different from the other cancers.

**Supplementary Figure 2:**
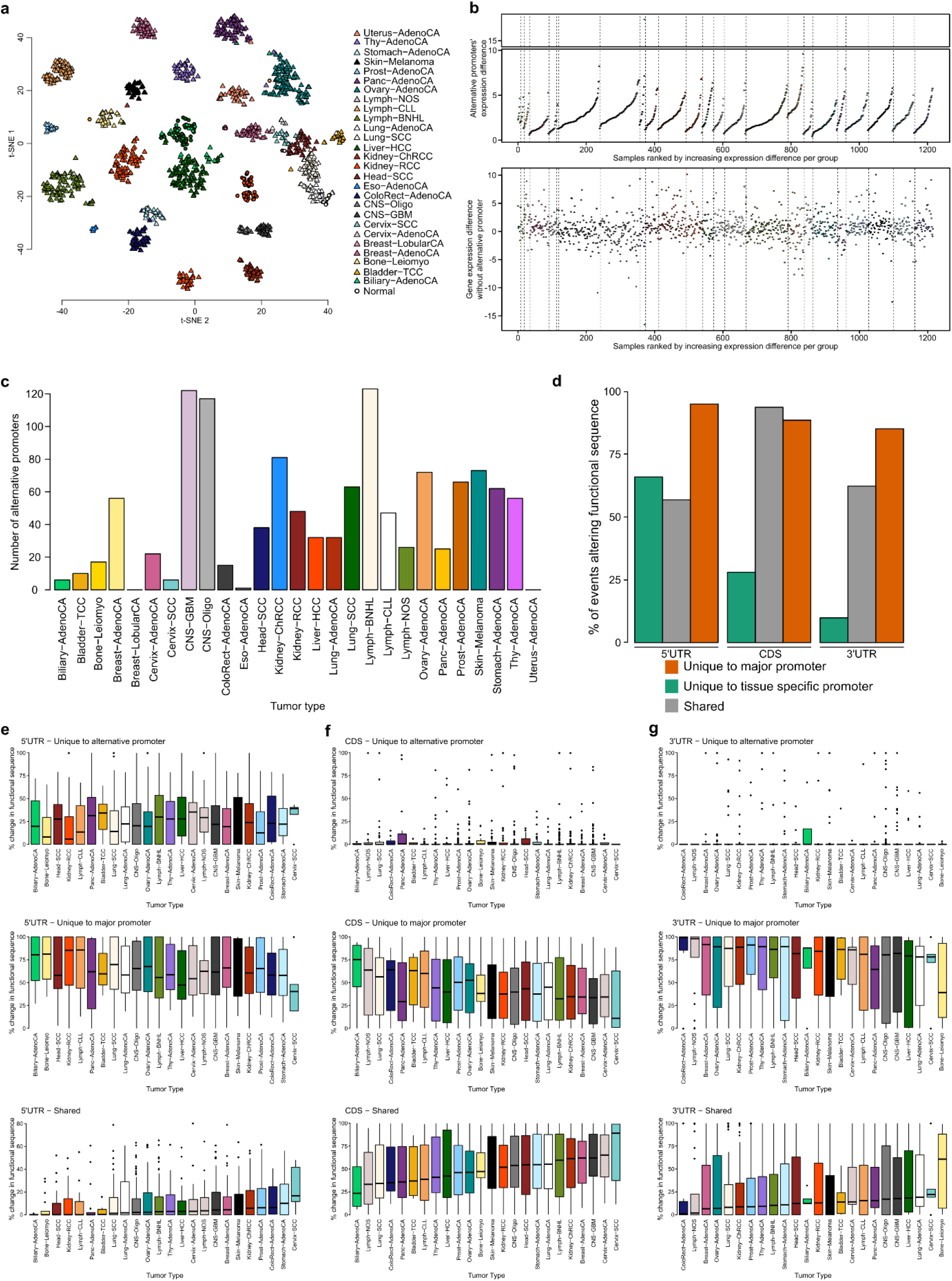
Alternative promoters display context specific regulation independent from the gene expression. (**a**) t-SNE plot using the top 1,500 genes with the highest variance in gene expression. (**b**) Difference in alternative promoter activities (upper panel) and gene expression excluding alternative promoters (lower panel) across pan-cancer. Alternative promoters’ contribution to tissue specificity is independent from gene expression. (**c**) Number of alternative promoters for each tumor type. (**d**) Shown is the percentage of times a change has occurred in the 5’UTR, 3’UTR, and CDS sequence that is unique to the tissue specific promoter (green), unique to the major promoter (orange), and that is shared (gray). (e, f, g) Shown is the percentage of the 5’UTR (**e**), CDS (**f**), and 3’ UTR (**g**) sequence that is unique to the tissue specific promoter gene isoforms (top panel), unique to the major promoter gene isoforms (middle panel), and that is shared (bottom panel).

**Supplementary Figure 3:**
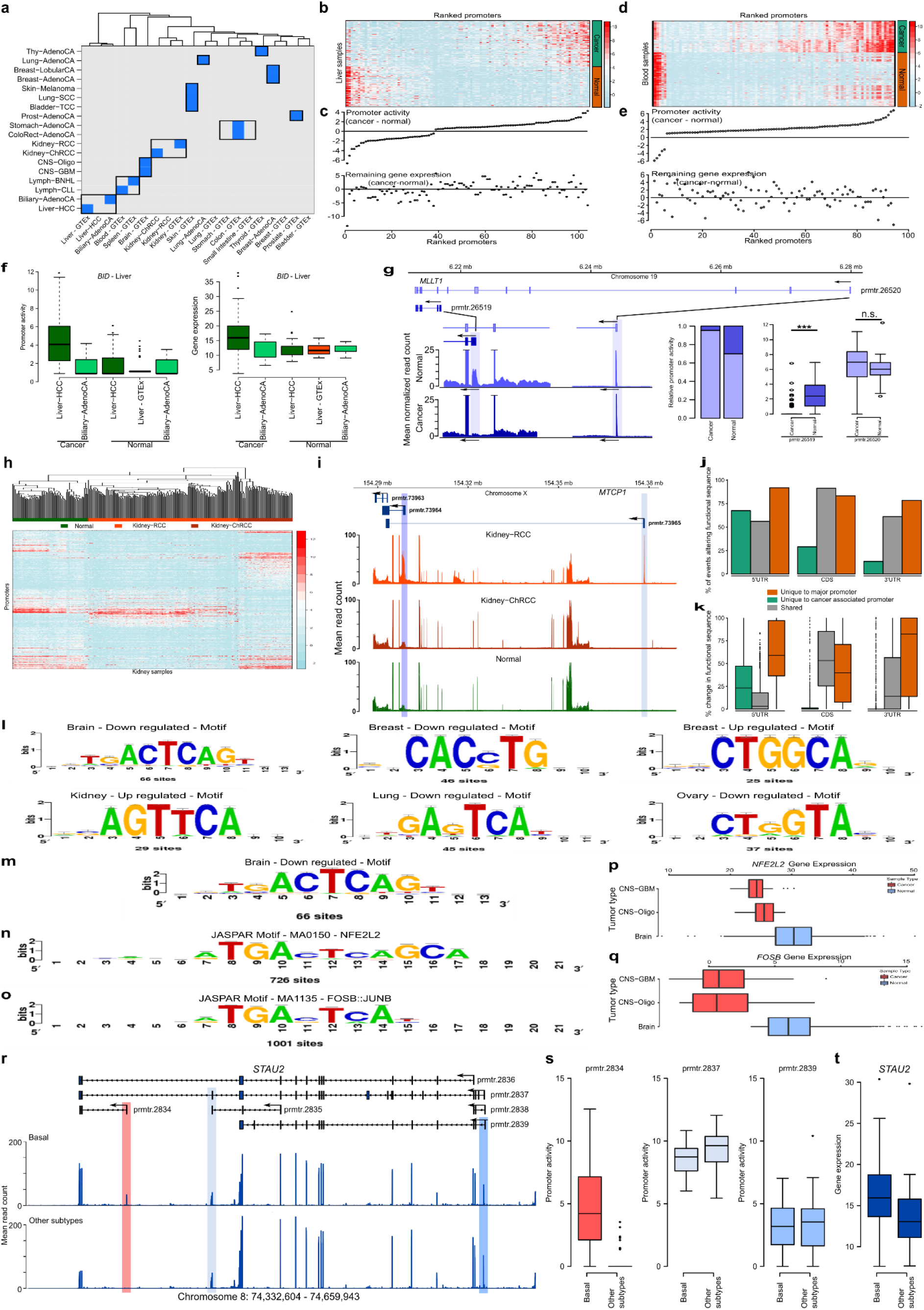
Identification of cancer-associated promoters. (**a**) Cancer samples from PCAWG match to normal samples from the same tissue (blue) regardless of data source (PCAWG or GTEx). Squamous cell carcinoma of the lung (Lung-SCC), and bladder cancer (Bladder-TCC) are assigned to skin reflecting the origin of cancer cell not the tissue. (**b**) Heatmap showing the normalized promoter activity estimates for liver cancer and normal samples, ranked by mean difference. (**c**) Difference in cancer-associated promoter activities (upper panel) and gene expression excluding alternative promoters (lower panel) for liver cancer and normal samples, ranking is similar to heatmap in (**b**). (**d**) Heatmap showing the normalized promoter activity estimates for blood cancer and normal samples, ranked by mean difference. (**e**) Difference in cancer-associated promoter activities (upper panel) and gene expression excluding alternative promoters (lower panel) for blood cancer and normal samples, ranking is similar to heatmap in (**d**). (**f**) Promoter activity and gene expression plots for the *BID* gene’s cancer associated promoter across liver cancer and normal samples. (**g**) Shown is the mean read count at the *MLLT1* gene locus for kidney tumor samples and normal samples (bottom-left). The light blue regions highlight the cancer-associated promoters in kidney cancer samples. The cancer associated deactivation of prmtr.26519 can be seen in relative (bottom-middle) and absolute (bottom right) promoter activities across normal and cancer samples. (**h**) Kidney cancer associated promoters display tumor type specific. (**i**) Shown is the mean read count for the *MTCP1* gene’s promoter that displays kidney cancer tumor type specific regulation. The first promoter (prmtr.73965) is only active in Kidney-RCC but not in Kidney-ChRCC tumors. (**j, k**) Shown is the percentage change in occurrence (**j**) and length (**k**) of the 5’UTR, 3’UTR, and CDS sequence that is unique to the cancer associated promoter (green), unique to the major promoter (**blue**), and that is shared (gray). (**l**) Sample motifs identified by RSAT de-novo motif discovery webserver to be enriched for different cancer associated promoter sets. (**m**) The PWM of the motif enriched for cancer associated down-regulated alternative promoters for brain tissue. (**n, p**) The transcription factor NFE2L2 binds to a similar motif to the identified de-novo motif for cancer associated down-regulated alternative promoters for brain tissue. (**n**) The JASPAR motif (MA0150) for NFE2L2 transcription factor. (**p**) The expression of NFE2L2 gene in brain cancer and normal samples. (**o, q**) The transcription factor FOSB::JUNB binds to a similar motif to the identified de-novo motif for cancer associated down-regulated alternative promoters for brain tissue. (**o**) The JASPAR motif (MA1135) for FOSB::JUNB transcription factor. (**q**) The expression of FOSB gene in brain cancer and normal samples. (**r**) Shown is the mean read count at the STAU2 gene locus for samples from the basal subtype of breast cancer and from all other subtypes of breast cancer. The light red, light blue and blue rectangle regions highlight the top 3 most active promoters. (**s**) The last STAU2 promoter (prmtr.2834) is more active in basal subtype compared to all other subtypes, whereas the other 2 active promoters (prmtr.2827 and prmtr.2839) show comparable activity levels in all subtypes. (**t**) Comparison of gene expression levels for STAU2.

**Supplementary Figure 4:**
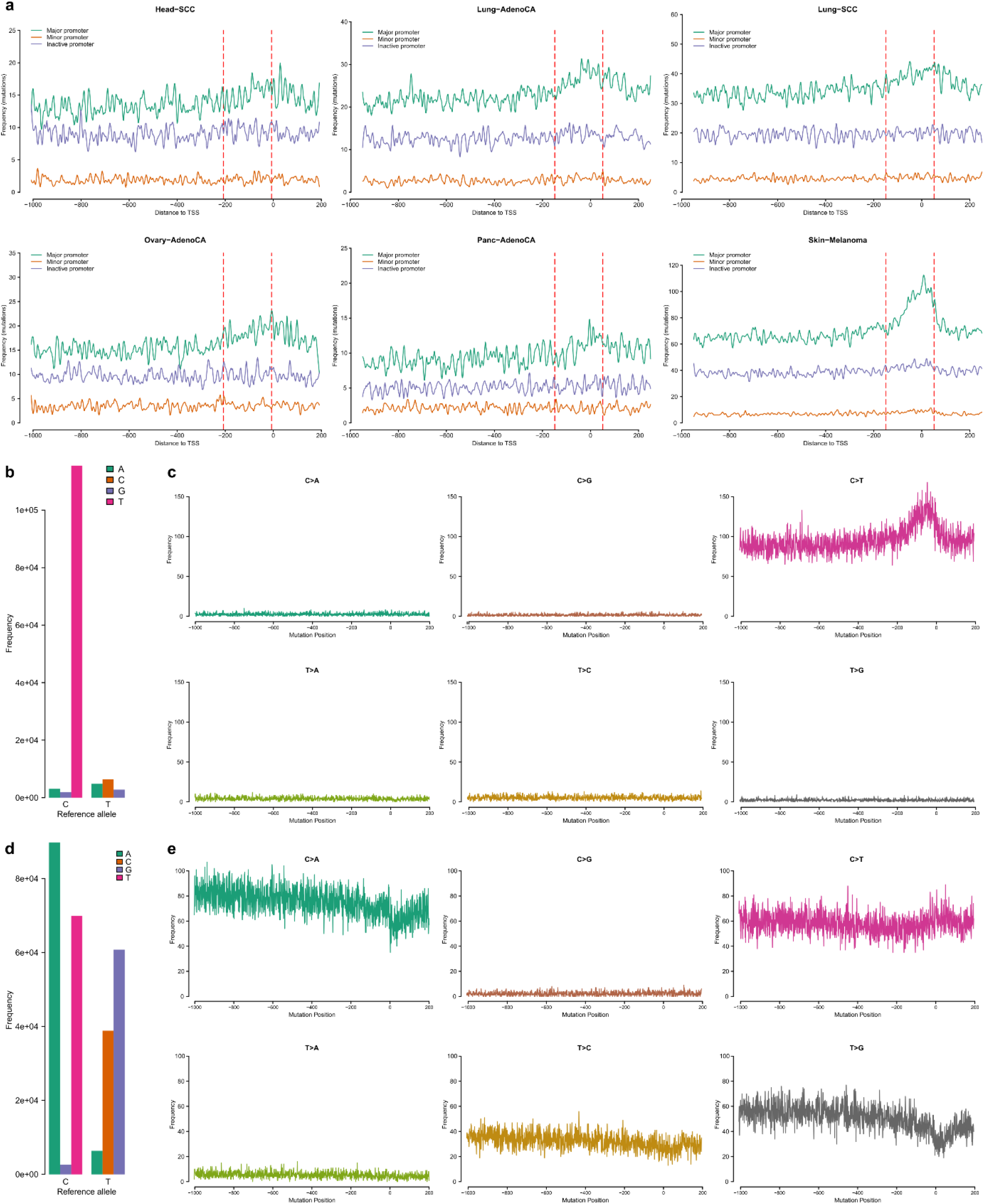
Overview of noncoding promoter mutation patterns across different cancer types. (**a**) Distribution of promoter mutations around major, minor and inactive promoters across cancer types where NER plays a major role. Red lines indicate the window of 200bps upstream TSS where major promoters show an enrichment of mutations whereas minor and inactive promoters don’t. (**b, c**) Overview of promoter mutations for Skin-Melanoma tumors. The majority of promoter mutations are C>T indicative of UV induced DNA damage (**b**). Distribution of promoter mutations for each mutation class reveals the enrichment of C>T mutations around the 200bps window upstream (**c**). (**d, e**) Overview of promoter mutations for ColoRect-AdenoCA tumors. The majority of promoter mutations are C>A and C>T (**d**). Distribution of promoter mutations for each mutation class doesn’t display an enrichment of mutations around the 200 bps window upstream differing from the mutation pattern of Skin-Melanoma tumors.

